# Selection on visual opsin genes in diurnal Neotropical frogs and loss of the *SWS2* opsin in poison frogs

**DOI:** 10.1101/2022.10.18.510514

**Authors:** YC Wan, MJ Navarrete, LA O’Connell, LH Uricchio, AB Roland, ME Maan, SR Ron, M Betancourth-Cundar, MR Pie, KA Howell, CL Richards-Zawacki, ME Cummings, DC Cannatella, JC Santos, RD Tarvin

## Abstract

Amphibians are ideal for studying visual system evolution because their biphasic (aquatic and terrestrial) life history and ecological diversity expose them to a broad range of visual conditions. Here we evaluate signatures of selection on visual opsin genes across Neotropical anurans and focus on three diurnal clades that are well-known for the concurrence of conspicuous colors and chemical defense (i.e., aposematism): poison frogs (Dendrobatidae), Harlequin toads (Bufonidae: *Atelopus*), and pumpkin toadlets (Brachycephalidae: *Brachycephalus*). We found evidence of positive selection on 44 amino acid sites in *LWS, SWS1, SWS2,* and *RH1* opsin genes, of which one in *LWS* and two in *RH1* have been previously identified as spectral tuning sites in other vertebrates. Given that anurans have mostly nocturnal habits, the patterns of selection revealed new sites that might be important in spectral tuning for frogs, potentially for adaptation to diurnal habits and for color-based intraspecific communication. Furthermore, we provide evidence that *SWS2*, normally expressed in rod cells in amphibians, has likely been lost in the ancestor of Dendrobatidae, suggesting that under low-light levels, dendrobatids have inferior wavelength discrimination compared to other frogs. This loss might follow the origin of diurnal activity in dendrobatids and could have implications for their chemical ecology, biodiversity, and behavior. Our analyses show that assessments of opsin diversification in understudied groups could expand our understanding of the role of sensory system evolution in ecological adaptation.

## Introduction

Natural selection has favored visual systems that maximize the detection and discrimination of light wavelengths that are relevant to organismal activity patterns (Warrant and Johnsen 2013). For instance, nocturnal or crepuscular species have visual systems that capture maximal light or permit color discrimination in dim-light conditions (Bowmaker 2008; Gutierrez et al. 2018; Mohun and Davies 2019; Guo et al. 2023). In contrast, diurnal species are active in bright and broad-spectrum light (e.g., 350-700 nm); therefore, their visual systems may be tuned for improved wavelength discrimination (i.e., better color vision; Bowmaker 2008). In vertebrates, visual opsin genes have diversified and undergone selection for adaptation to various environments. For example, diurnal primates underwent duplication and spectral tuning of the long-wavelength-sensitive opsin twice (Hunt et al. 1998) and fishes have undergone duplication and diversification of opsin genes many times (Hofmann and Carleton 2009). In addition, some groups have lost opsin genes during adaptation to new light environments including the loss of cone opsins in deep-sea coelacanths (Yokoyama et al. 1999) and loss of a short-wavelength sensitive opsin in mammals during adaptation to nocturnality (Bowmaker 2008).

Vision in anuran amphibians is particularly interesting as the visual system often needs to transition from functioning in an aquatic environment during the larval stage to a terrestrial environment during the juvenile phase. Each setting presents unique visual requirements (Cronin et al. 2014). Furthermore, while most frogs are primarily nocturnal or crepuscular, a few clades have evolved diurnal activity (Anderson and Wiens 2017). Due to the selective pressure of diurnal and visually guided predators, many diurnal frogs are adorned with colorful visual signals that evolved as part of a defensive strategy known as aposematism, in which conspicuous signals warn predators of prey defenses (Bell and Zamudio 2012). Although aposematic signals initially evolve under selective pressure from predators and might intensify over evolutionary time (Mappes et al. 2005; Sherratt 2008; Loeffler-Henry et al. 2023), warning signals can also become entangled with mating behaviors and further diversify under sexual selection pressures (Cummings and Crothers 2013; Rojas et al. 2018), if the mate recognition and visual sensory system is appropriately tuned to the signal.

The molecular underpinnings of vision in vertebrates rely on a set of photosensitive opsin genes that encode diverse G-protein-coupled receptors that bind to a retinal chromophore (Bowmaker 2008). The visual opsin genes present in vertebrates include long-wavelength sensitive (*LWS* or *OPN1LW*), middle-wavelength sensitive (*RH2, MWS* or *RHB*; not to be confused with an independent gene duplication that gave rise to a middle-wavelength-sensitive opsin gene in Old World primates also called *MWS,* or *OPN1MW* [Nathans et al. 1986; Hunt et al. 1998]), short-wavelength sensitive 1 (*SWS1* or *OPN1SW*), and short-wavelength sensitive 2 (*SWS2* or *OPN2SW*; Bowmaker 2008; Schott et al. 2022). Other vision-related genes include the rod opsin, rhodopsin (*RH1*, *RHO, RHA,* or *OPN2*), which evolved after a duplication and divergence event from an ancestral opsin shared with *RH2* (Okano et al. 1992). Color vision is achieved when two or more photoreceptors with opsins differing in wavelength sensitivity are simultaneously activated, and their signals are compared by the observer’s central nervous system (Bowmaker 2008; Gibson 2014). For consistency, we refer to the opsin genes using the following acronyms: *LWS, SWS1, SWS1, RH1, RH2*.

The anuran retina is thought to typically contain three types of cone cells that are active under bright-light conditions: two types of long-wavelength-sensitive cones that express *LWS*, and a short-wavelength-sensitive cone that expresses *SWS1* (Liebman and Entine 1968; Hárosi 1982; Koskelainen et al. 1994; Mohun and Davies 2019; Donner and Yovanovich 2020). However, some anuran species such as *Rana pipiens* (nocturnal) and *Oophaga pumilio* (diurnal), possess an additional fourth cone with middle-wavelength sensitivity (λ_max_ of ∼500 nm), but the pigment contained in this type of photoreceptor cell remains to be determined (Siddiqi et al. 2004; Mohun and Davies 2019). In some vertebrates (e.g., fish and reptiles; Bowmaker 2008), the *RH2* gene is responsible for green or middle-wavelength sensitivity. To date, the *RH2* gene has not been found in any amphibian, and it is believed to have been lost in their last common ancestor (Mohun and Davies 2019; Schott et al. 2022). Thus, it is plausible that *RH1* is the gene that is expressed in green-sensitive cones in amphibians (Schott et al. 2022), conferring them with the ability to detect middle-wavelength light spectra during the day, as has occurred in at least one species of snake (Schott et al. 2016). In addition to these four cone cell types (single and double LWS cones, single SWS1 cones, and an MWS cone), the anuran retina has two types of rod cells that are active in dim-light conditions: one that expresses *RH1*, known as the "red rod," and another that expresses *SWS2*, known as the "green rod" (Hunt and Collin 2014). The existence of two rod types likely allows some amphibians to have low-light (scotopic) and rod-based color discrimination, but this has not been empirically tested (Yovanovich et al. 2017). Although some vertebrates use oil droplets to further tune wavelength sensitivity, oil droplets in anurans are often absent or colorless and thus are unlikely to influence the spectral sensitivity of cone opsin proteins (Toomey and Corbo 2017). For example, oil droplets are present but colorless in *O. pumilio* (Siddiqi et al. 2004).

While a few anuran models (e.g., *Rana temporaria, Xenopus laevis,* and *Rhinella marina* [*Bufo marinus*]) have played critical roles in discovering the biology of vision, none of these species are diurnal and the vast diversity of anuran visual systems is only just beginning to be unraveled (Donner and Yovanovich 2020). For example, only 108 of the >7500 currently described anuran taxa (AmphibiaWeb 2022) have information on lens transmission properties (Yovanovich et al. 2020; Thomas et al. 2022), and the spectral sensitivity of retinal photoreceptor cells has been described for only 10 species (Donner and Yovanovich 2020). Moreover, the genetic basis of anuran vision is especially understudied (Mohun and Davies 2019), with few genomic resources until very recently limited to a handful of well-assembled frog genomes (Womack et al. 2022) and one recent study of opsin genes in 33 frogs (Schott et al. 2022).

If compared to other vertebrates, this lack of information on the physiology of anuran vision contrasts with the relatively large body of research into anuran visual ecology (e.g., Toledo and Haddad 2009; Bell and Zamudio 2012; Rößler et al. 2019). One example is Dendrobatidae (poison frogs), which has become a model system for understanding how natural selection by predators shapes the origin and subsequent diversification of aposematic signals (e.g., Clough and Summers 2000; Richards-Zawacki and Cummings 2011; Santos and Cannatella 2011; Wang 2011; Cummings and Crothers 2013; Rojas et al. 2014; Lawrence et al. 2019). Dendrobatid warning signals can also play a role in intraspecific communication. For instance, color perception seems to be key for mate recognition and territorial display. In the aposematic species *O. pumilio,* tadpoles imprint on their mother’s color (Yang et al. 2019), females show assortative mate preferences (Summers et al. 1999; Reynolds and Fitzpatrick 2007; Maan and Cummings 2008; Yang et al. 2016), male-male competition relies on color-mediated aggressive behaviors towards phenotypically similar rivals (Yang et al. 2018; Yang and Richards-Zawacki 2020), and there may be directional sexual selection on male coloration brightness (Maan and Cummings 2009).

Assortative mating by coloration and pattern is present but less pronounced in other dendrobatid species of the same genus (*O. histrionica* and *O. lehmanni*; Medina et al. 2013) and sexual dichromatism in *Mannophryne* where throat coloration in males (gray) and females (bright yellow) plays a role in territoriality and mate choice (Greener et al. 2020). Aposematic dendrobatids are also a special focus of molecular work, including two public genomes (i.e., *Oophaga pumilio* and *Ranitomeya imitator*), with several more being assembled.

Other frog families that include diurnal and aposematic species include some Neotropical genera of Bufonidae and Brachycephalidae. *Atelopus* (Harlequin toads, Bufonidae) and *Brachycephalus* (Pumpkin toadlets, Brachycephalidae) have a handful of studies on color-based intraspecific or visual signaling in brightly colored species. For example, *Atelopus zeteki* appears to use limb motions as visual signals to conspecifics and this might be related to their lack of tympanic middle ear (Lindquist et al. 1998); yet it is unknown if other *Atelopus* use their conspicuously colored soles for intraspecific communication (Rößler et al. 2019). Likewise, at least one species of *Brachycephalus* is inferred to be aposematic based on experimental evidence (Goutte et al. 2019; Rebouças et al. 2019), and some species of *Atelopus* have conspicuously colored soles that might be aposematic signals (Rößler et al. 2019); *B. ephippium* and *B. pitanga* also have fluorescent dermal bones visible through their skins with a potential function as a signal (Goutte et al. 2019). While territorial males of some *Brachycephalus* display a foot-waving behavior to warn off encroaching males similar to some species of *Atelopus* (Pombal et al. 1994), it is unclear how important a role color and/or contrast play in the display. To our knowledge, there have not been any studies examining color-based mate preference in these genera. Lastly, no genomic or transcriptomic work focusing on genes involved in vision in *Atelopus* and *Brachycephalus* have been conducted, and no publicly available genomes from either group exist.

Here we aimed to describe patterns of selection on visual opsin genes in diurnal and three colorful frog lineages of the Neotropics. We sequenced the four amphibian visual opsin genes known in amphibians (*LWS*, *SWS1*, *SWS2* and *RH1*) using a target-bait capture approach. Our taxon sampling included 116 species with a special emphasis on the diurnal frog clades: poison frogs (Dendrobatidae or Dendrobatoidea sensu Grant *et al*., 2006), Harlequin toads (Bufonidae: *Atelopus*), and pumpkin toadlets (Brachycephalidae: *Brachycephalus*). We reviewed amino acid sites that are relevant to the spectral sensitivity of opsin proteins, which may have been modified by natural selection via substitutions to better absorb wavelengths that are most relevant to the organism’s visual ecology, a process known as spectral tuning (Carvalho et al. 2007; Hunt et al. 2007; Osorio and Vorobyev 2008). Using a combination of site and branch-site selection analyses, we determined whether amino acid positions at or near known spectral tuning sites were under selection, and whether these substitution patterns differed between diurnal and nocturnal lineages. In regards to the *MWS* cone found in some amphibians, our results are concordant with others (Schott et al. 2022), and propose a potential role for *RH1*. Additionally, we were unable to capture sequences of *SWS2* from any poison frog species, which was further confirmed with transcriptome sequencing and genome canvassing that suggest the absence of this gene in Dendrobatidae. This gene was found in other frogs, which supports a hypothesis that poison frogs have lost *SWS2,* likely in their last common ancestor between ∼70 and 40 million years ago (Mya).

## Results

### Opsin sequences and phylogenies

We retrieved 28 opsin gene sequences of 13 species from GenBank and reconstructed another 49 (representing 19 species) using SRA data deposited in NCBI (Table S1). From 484,530 Illumina reads from our bait capture experiment, we further reconstructed 289 new opsin gene sequences representing 87 anuran species. In total we analyzed 366 sequences from 116 anuran species. All assembled sequences had a read coverage of >10x, with 82% having coverage >50x (Table S2). Estimated gene trees were largely concordant with recent family-level phylogenies (Feng et al. 2017; Hime et al. 2020) and hypothesized interspecific relationships (Pyron 2014; Grant et al. 2017; Jetz and Pyron 2018; see Data S3). The phylogeny inferred using all opsin gene sequences was reciprocally monophyletic for each gene, indicating a low probability of chimeric sequences in our dataset.

**Table 1.**
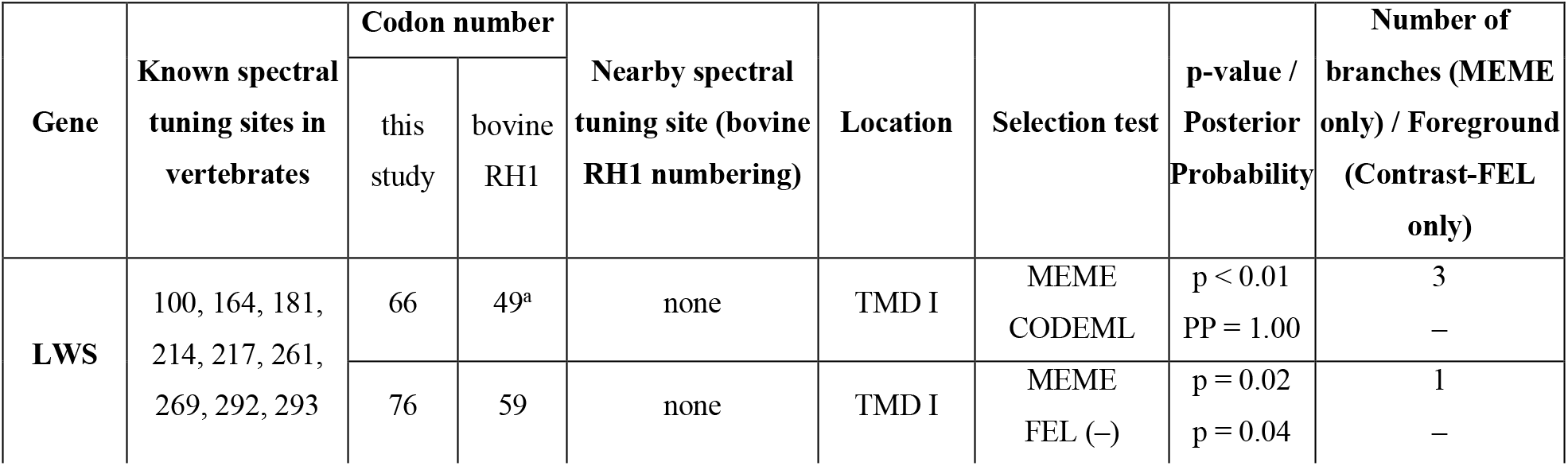
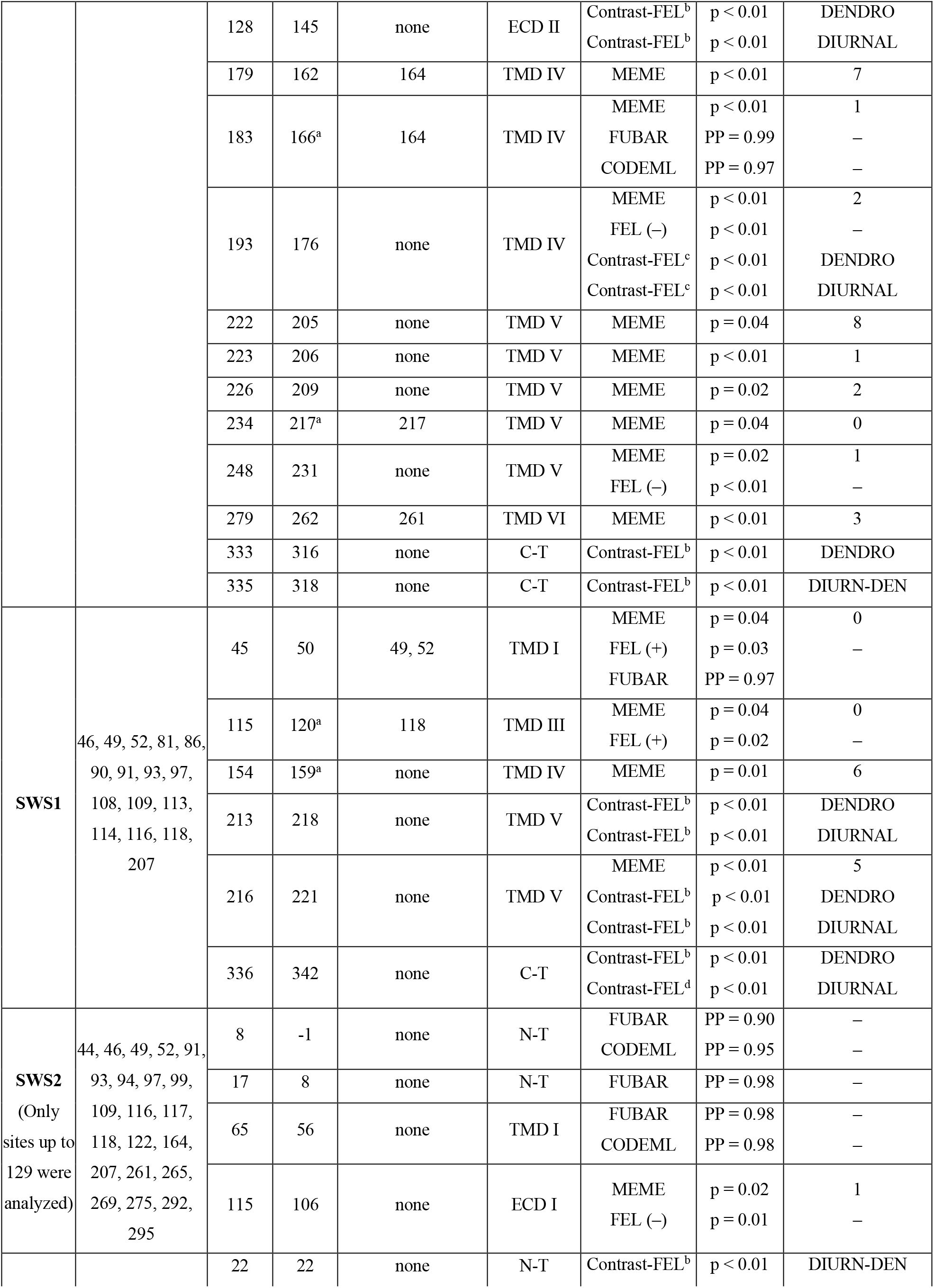
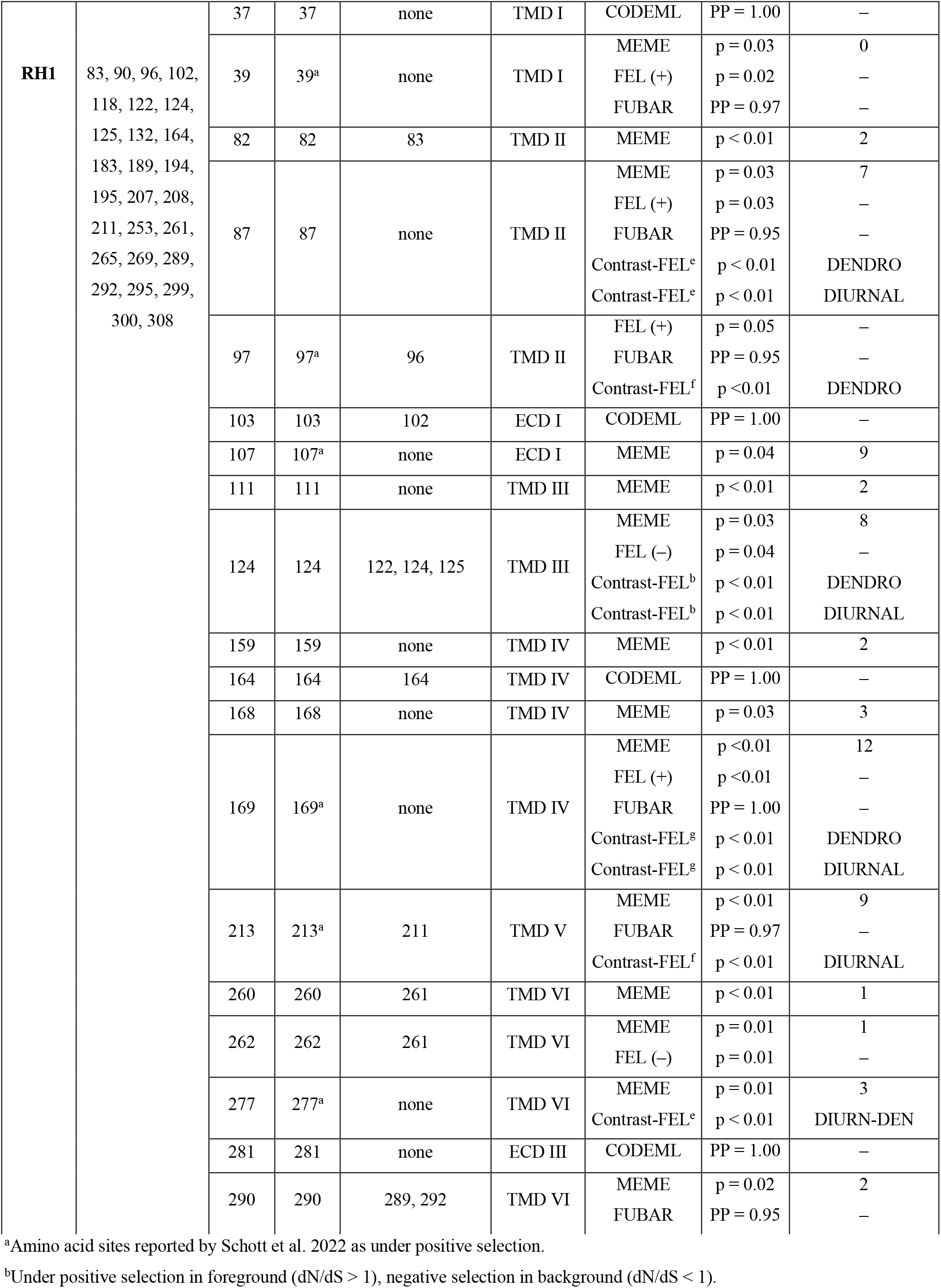
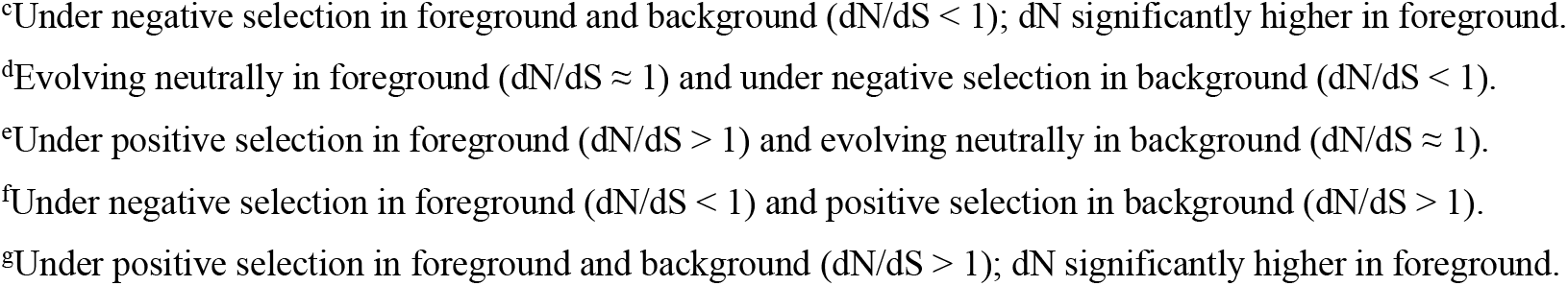
Selection analysis results. Amino acid sites in four opsin genes that were detected to be under positive selection by HyPhy and CODEML analyses. We aligned sequences against bovine rhodopsin (NP_001014890.1) to make comparisons with known spectral tuning sites (see Table S3 for references). More details regarding the phylogenetic patterns of selection can be reviewed in Fig. 1 and S1. We report all significant FEL results that correspond to sites identified by MEME or FUBAR, and note each FEL result as indicative of positive (+) or negative (–) selection. CODEML analyses are reported as posterior probabilities comparing M7 and M8 models with Bayes Empirical Bayes analysis. Foreground groups for Contrast-FEL are indicated by “DENDRO”, i.e., Dendrobatidae + stem branch, “DIURNAL”, i.e., all diurnal branches as described in methods, or DIURN-DEN, which is all diurnal branches except for Dendrobatidae. N-T, N-terminus; C-T, C-terminus; ECD I, extracellular domain I; TMD I–VII, transmembrane domains I–VII; –, does not apply. Reported p-values for MEME have been corrected for multiple testing using Holm-Bonferroni; for Contrast-FEL, we report only the sites that pass the False Discovery Threshold (q = 0.02). Input files and results are available in Data S1.

### Analyses of selection

CODEML and HyPhy selection tests identified a total of 44 sites across the four opsin genes as having experienced positive selection (Table 1). Of these sites, 14 were identified in *LWS*, 6 in *SWS1*, 4 in *SWS2,* and 20 in *RH1*. We used the bovine sequence to identify site position and compare to prior empirical data (see Table S3); based on these data, 3 of the 44 sites are known spectral tuning sites including site 217 in *LWS* and sites 124 and 164 in *RH1,* and 12 other sites are within three amino acids from a known spectral tuning site (Table 1; Data S1; Table S3). In 11 of the 44 identified sites, our results agree with Schott and colleagues (2022) as being under positive selection (see superscripts in Table 1). Across the phylogeny, most sites were found to be under positive selection in only one or two branches, but 13 (sites 49, 162, 205, and 262 in *LWS,* sites 159 and 221 in *SWS1,* and sites 87, 107, 124, 168, 169, 213, and 277 in *RH1*) were found to be under positive selection in three or more branches, suggesting that these sites may be of particular functional importance in frogs (Figs. 1 and S1).

**Figure 1.**
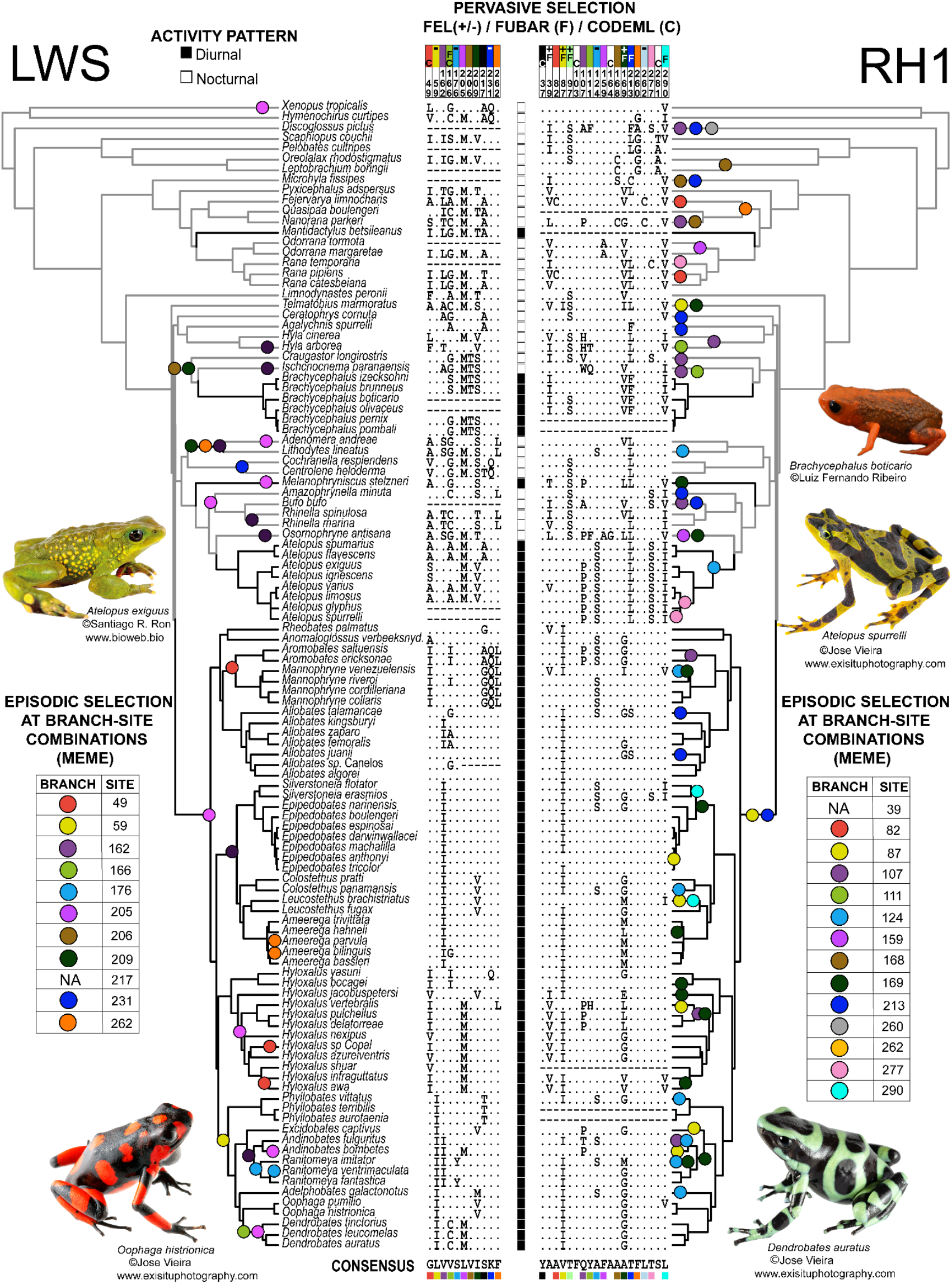
A summary of activity trait data (see Table S1 for references) and results from HyPhy and CODEML selection analyses of two opsin genes, *LWS* and *RH1* (see Fig. S1 for *SWS1* and *SWS2* and Table 1 for details) using bovine *RH1* numbering. Branch-site combinations that were detected by MEME (Mixed Effect Models of Evolution) as evolving under positive selection are mapped onto the phylogenetic tree using filled circles where circle colors correspond to specific sites in the alignment. One site in *LWS* and one in *RH1* were significant in MEME analyses but no specific branch-site combination was reported; these are marked by “NA” in the MEME tables. Amino acid sites marked with “F” or “C” above the alignment were reported to be under positive selection by FUBAR or CODEML, respectively. FEL results are indicated using + to indicate positive selection and – to indicate negative selection (negative selection results are only shown for sites found to be under positive selection by other analyses; see Data S1 for full FEL results). We abbreviated *Anomaloglossus verbeeksnyderorum* as *Anomaloglossus verbeeksnyd.* for brevity. Black branches indicate branches categorized as diurnal.

Among the list of sites in Table 1, some present complex characterizations. These include site 277 in *RH1*, which was reported to be under positive selection by MEME. However, we noted that there were no non-synonymous changes on the branches identified by MEME (*Atelopus spurrelli* and Node 33, the ancestral branch leading to *Atelopus spurrelli, A. glyphus, A. limosus,* and *A. varius*; Fig. 1). Instead, these branches have two mutations that do not result in an amino acid substitution (Ser-AGC → Ser-TCC), but likely require a nonsynonymous intermediate that is not recovered in our analysis.

Another six sites were found with contradictory results between analyses such that each site was detected by MEME to be under positive selection in a subset of branches but detected by FEL to be under negative selection across the entire phylogeny (Table 1, denoted with “FEL (–)”). Five of these sites are characterized by high conservation, with an amino acid change present in only one or two branches (Figs. 1 and S1). In contrast, the sixth site, site 124 in *RH1*, presents an alanine-to-serine replacement in eight branches. Finally, one site in *LWS*, two sites in *SWS1*, and one site in *RH1* (Table 1) were found to be under positive selection by MEME but without any branch identified, suggesting that the signal for selection is diffuse, which occurs when there is enough signal to report positive selection, but no individual branch rises to the level of significance (Spielman et al, 2019).

In order to identify patterns of selection associated with diurnality, we used Contrast-FEL to identify sites under differential dN/dS regimes between diurnal and nocturnal clades (“DIURNAL” foreground, see methods and Table S1 for how we determined activity states) on each gene tree except *SWS2,* for which no sequences from Dendrobatidae were available. This analysis identified two sites in *LWS,* three sites in *SWS1,* and four sites in *RH1* to be under differential selection regimes in foreground and background lineages (Table 1). Because all these sites had been identified by other methods as under positive selection when the entire tree was included (using FEL, CODEML, FUBAR, or MEME approaches), we suspected that our large sampling of dendrobatid lineages (all diurnal) might be driving this pattern. We then analyzed the dN/dS rates in Dendrobatidae branches plus its stem branch (“DENDRO” foreground, see Table 1) versus other branches. This analysis identified 7 of the 9 sites found by the DIURNAL analysis. In addition, one site was found to be evolving under significantly different regimes in the DIURNAL foreground and background but was not identified in the DENDRO analyses (site 213 in *RH1*). Interestingly this site was identified to be under negative selection in the foreground and under positive selection in the background, suggesting that it might be under selection in nocturnal lineages. A second site (site 342 in *SWS1*) was found to be under positive selection in the foreground, but under negative selection in the background for DENDRO; when setting the DIURNAL lineages to foreground, this site was found to be neutrally evolving in the foreground and under negative selection in the background. This result suggests that including additional diurnal lineages diluted a Dendrobatidae-specific pattern. In addition, two sites were found to be evolving under different regimes in the DENDRO foreground and background but were not identified in the DIURNAL analyses (site 318 in *LWS* and site 97 in *RH1*), suggesting that these sites are uniquely under selection in Dendrobatidae.

To further assess whether the amino acid sites identified to be under selection in diurnal lineages were driven by our biased sampling of Dendrobatidae and to potentially identify amino acid sites under selection in non-dendrobatid diurnal lineages, we then created a third group (“DIURN-DEN”), which is a subset of the DIURNAL group excluding Dendrobatidae and its stem branch. In these analyses, three sites were identified to be under differential selective regimes (318 in *LWS* and 22 and 277 in *RH1*) and they did not overlap with any sites identified to be under differential selective regimes when using the DIURNAL or DENDRO foregrounds. Each of these sites was found to be under positive selection in foreground and to be under negative selection or neutrally evolving in background, suggesting that they might be adaptations to diurnality in non-dendrobatid clades.

Based on these results, we note that caution should be taken when contrasting branches for selection pattern analyses based on a phylogeny that has multiple independent origins of a trait. The uneven distribution of taxa (e.g., overrepresentation of clades with the derived character of interest–in our case Dendrobatidae) might drive the analyses to identify sites with a signature of selection exclusive to the overrepresented group as significant in overall selection analyses.

### *Verifying absence of* SWS2 *in poison frogs*

During our exome capture reconstructions, we failed to recover any exon or fragments of the *SWS2* gene in dendrobatids. We considered at first that our baiting design might have been unsuccessful, which would suggest that our *SWS2* sequence baits (designed using a consensus sequence matrix) were not similar enough to capture this gene, but we had no issues obtaining sequences from all other clades including the other aposematic species of *Atelopus* and *Brachycephalus* (Fig. S1). Therefore, we needed to confirm that this gene was not present with additional transcriptomic and genomic data. For the transcriptome approach, we extracted mRNA from the *O. pumilio* eye, and we were unable to amplify *SWS2* cDNA. This suggests that at least in this species, *SWS2* is not expressed. For the genomic approach, we used synteny analyses with BLAST and we found that *SWS2* is located in a syntenic block with *LWS* and *MECP2* in the ancestors of all tetrapods, all amniotes (Fig. 2A), and in five frog families with genomic data that is publicly accessible (Fig. 2B: Bufonidae, Leptodactylidae, Eleutherodactylidae, Pipidae, and Ranidae). For dendrobatids, we explored the reassembled *O. pumilio* scaffolds and found that scaffold70671 contained *LWS* and was syntenic with other amphibians (two short and highly conserved regions between *SWS2* and *LWS* could be aligned between *Nanorana* and *O. pumilio* [Data S2]), but we could not identify any coding region of *SWS2*. Then, we explored the more complete scaffold of this region from *R. imitator* (scaffold934), which also included *LWS* and the gene *MECP2*, the latter of which is expected to be upstream of *SWS2*, yet we were unable to find *SWS2* between these two genes on this scaffold (Fig. 2B). In fact, we could not detect an *SWS2* gene or pseudogene on any scaffold in *O. pumilio* or *R. imitator* genomes (Table S4). In contrast, using the same methods, we were able to detect *SWS2* upstream of *LWS* in other hyloid frogs including *Bufo, Eleutherodactylus*, *Engystomops*, and *Rhinella* (Fig. 2B, Data S4). The hyloid clade originated ∼70 Mya and the age of the last common ancestor of Dendrobatidae has been estimated to be ∼40 My (Feng et al. 2017). Thus, the most parsimonious explanation is that a functional *SWS2* gene is not present in living dendrobatids, including *O. pumilio* or *R. imitator*, and that the *SWS2* gene may have been pseudogenized (to a point beyond recognition) or lost in the ancestor of dendrobatids between 70 and 40 Mya.

**Figure 2.**
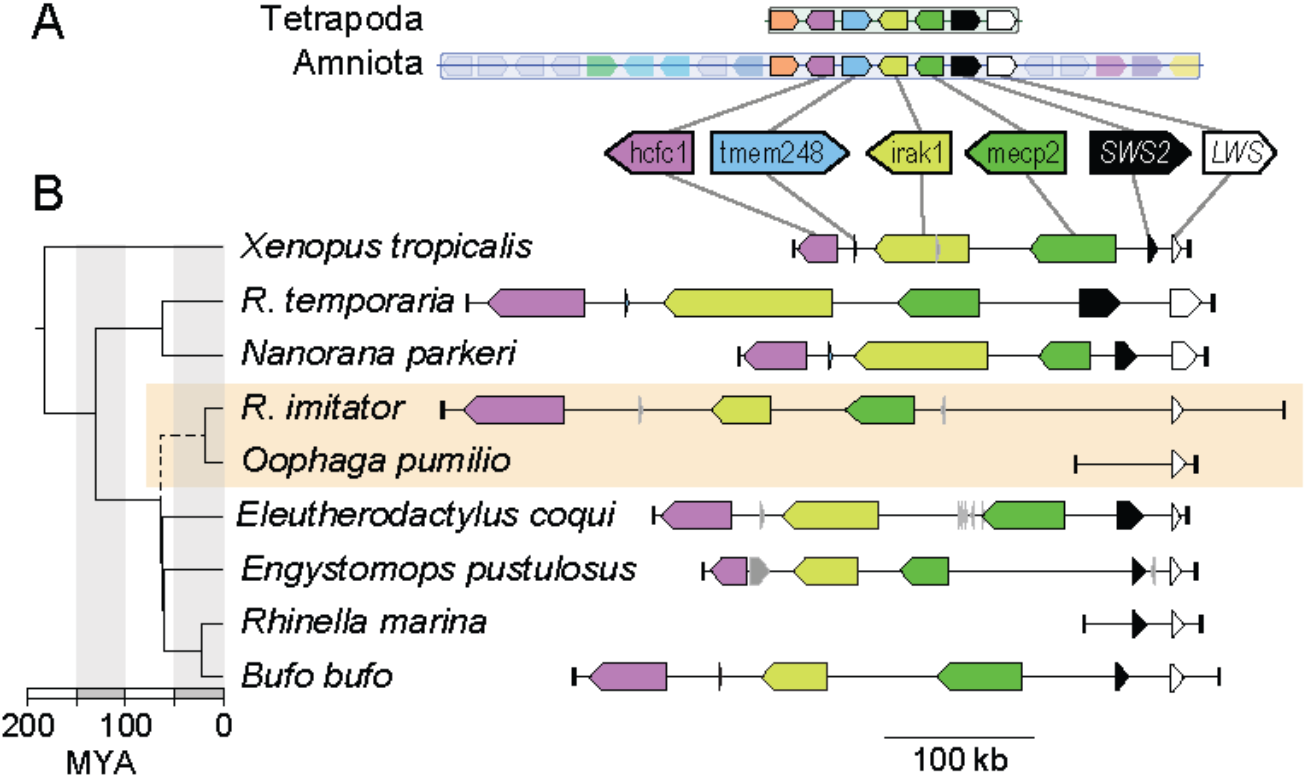
A) *SWS2* is found in a syntenic block with *LWS* in the ancestors of all tetrapods (block 355) and in the ancestor of all amniotes (block 65). B) The region in *Xenopus tropicalis* (chromosome 8, NC_030684:32108538- 32360324) shows synteny with *Rana (R.) temporaria* (chromosome 9, NC_053497.1:22641469-22170306; Ranidae), *Nanorana parkeri* (unplaced scaffold, NW_017306743:511492-806940; Ranidae), *Rhinella marina* (contig ctg22529_RHIMB, ONZH01019223.1:1-80000; Bufonidae), *Bufo bufo* (chromosome 8, NC_053396.1:25948637-26357822; Bufonidae), *Eleutherodactylus coqui* (chromosome 9, CM034094.1:11013102-11351932; Eleutherodactylidae), *Engystomops pustulosus* (chromosome 10, CM033650.1:68367373-68196643; Leptodactylidae), which also all contain *SWS2, MECP2, IRAK1, TMEM187,* and *HCFC1* upstream of *LWS*. However, *SWS2* is absent in the scaffold containing *LWS* in two dendrobatid genomes (*Oophaga pumilio* [scaffold70671:1-49101] and *Ranitomeya* [*R.*] *imitator* [CAJOBX010072427.1, scaffold934:531527-1]). A dated chronogram (Feng et al. 2017) indicates that *SWS2* may have been lost in the ancestral lineage (dashed line) leading to Dendrobatidae (orange box), between ∼70 and 40 Mya. Where coding regions were identified in genome assemblies but not annotated we added annotations using blastp and blastn. Coding regions in grey had ambiguous blast results and thus were left without annotations.

## Discussion

In this study, we reviewed the evolution of the visual opsin gene repertoire (*LWS*, *SWS1*, *SWS2*, and *RH1*) in anurans by analyzing a dataset of 116 frog species from 20 families, including nocturnal, diurnal, and crepuscular species. Our analyses report 44 amino acid sites as under positive selection across Anura, including sites in or near locations that have been implicated in visual tuning. Many of these sites are undergoing positive selection in branches leading to or within diurnal and aposematic Neotropical frog clades. Moreover, our results provide evidence for the loss of the *SWS2* gene in the ancestor of Dendrobatidae between 70–40 Mya, an event that coincides with the evolution of diurnal activity patterns in this clade. Diurnality in poison frogs has been assumed as a prerequisite for the origin of many of their unique adaptations including aposematism, audiovisual communication, mating choice, and parental care.

### Selection analyses

Most of the amino acid sites across the four opsin genes were under negative or purifying selection (Data S1). The sites inferred to be under positive selection occur predominantly in the transmembrane domains (Table 1). Transmembrane domains impact the tertiary structure, thermal stability, and properties of the retinal-binding pocket (Andrés et al. 2001; Yokoyama et al. 2006). Spectral sensitivity is related directly to interactions between the amino acid residues in the transmembrane domains and the chromophore (Yokoyama et al. 2006). Although it is possible that replacements at these sites might influence spectral tuning in frogs, without more experimental data, their implications are difficult to interpret. For this reason, we conservatively discuss their possible role in anuran vision below.

Of the 44 sites we found to be under positive selection, only three are previously known spectral tuning sites in vertebrates. A substitution from nonpolar alanine to polar serine at site 217 in *LWS* (233 in the human *LWS*) has been previously shown to play a role in the differentiation of the spectral sensitivities between the long-wave-sensitive and the middle-wave-sensitive pigments in humans and other mammals (Yokoyama and Yokoyama 1990; Asenjo et al. 1994; Fasick and Robinson 1998; Yokoyama and Radlwimmer 2008, but see Hiramatsu et al. 2004). In our study, MEME detected positive selection on this site without indicating any specific branches, suggesting a diffuse signal of selection at this site across the tree (Fig. 1; Table 1). Moreover, this site was also reported by Schott et al. (2022) to be under positive selection, confirming that this is a key site that could be responsible for shifting the sensitivity of *LWS* in frogs, including the diurnal taxa explored in this study. In our alignment, amino acid substitutions at site 217 are present in several species including several diurnal and colorful clades such as the dendrobatid genera *Mannophryne, Rheobates, Aromobates,* and *Phyllobates*, as well as two species of *Atelopus*. However, this site was not identified in Contrast-FEL analyses as under positive selection specifically in diurnal lineages.

Another known spectral tuning site in RH1 (site 124) was found to be under positive selection by MEME in eight branches (Ranitomeya imitator, Adelphobates galactonotus, Andinobates fulguritus, Phyllobates vittatus, Colostethus panamansis, Lithodytes lineatus, Mannophryne venezuelensis, and the branch leading to the Atelopus clade). The amino acid type at site 124 of those species is serine instead of alanine as seen in other anuran species. Contrast-FEL analyses found this site to be under positive selection in diurnal lineages and in dendrobatid lineages but under negative selection in other branches. Mutagenesis experiments in bovine RH1 have shown that alanine-to-threonine and alanine-to-serine substitutions at this position resulted in a slight blue-shift (Lin et al. 1998; Castiglione and Chang 2018). A124S in combination with L119F significantly increases the stability of the active-conformation of bovine RH1 (Castiglione and Chang 2018). Therefore, it is possible that an anuran RH1 with the A124S substitution has a blue-shifted λ_max_, and that such changes at this site may be associated with adaptation to diurnality as has been suggested in species of diurnal snakes (Schott et al. 2016; Hauzman et al. 2017).

In addition, site 164 in *RH1*, another known spectral tuning site, was found by our CODEML analysis to be under positive selection; other analyses did not identify this site to be under selection. In our dataset, 102 species at this position had an alanine, while just *Osornophryne antisana* expressed a glycine. Substitutions at this site are known to contribute to a red shift in the absorption maxima of bovine *RH1* (Chant et al. 1992). Although an A164S substitution seems to result in a rather small (2 nm) red shift in absorption, it has been shown that an additive effect is achieved when it is in combination with other substitutions including F261S and A269T (Chan et al. 1992). Thus, it is difficult to predict how changes at this site might impact the visual acuity of *O. antisana* or other frog species, and our analyses do not offer strong support for the involvement of site 164 in *RH1* in adaptation to diurnality.

Considering the general lack of data on spectral tuning sites in frog opsins, it is possible that other sites we identified to be under positive selection (Table 1) could be directly or indirectly involved in spectral tuning in anurans. Eleven sites found to be under positive selection are not known spectral tuning sites but are located within three amino acids of a known spectral tuning site: 162, 166, and 262 in *LWS*; 50 and 120 in *SWS1*; 82, 97, 103, 213, 260, 262, and 290 in *RH1*. Additionally, thirteen sites, including three at or near spectral tuning sites, were found to be under selection in three or more branches:: 49, 162, 205, and 262 in *LWS*; 159 and 221 in *SWS1*; 87, 107, 124, 168, 169, 213, and 277 in *RH1* (Table 1, Fig. 1). We speculate that some of these sites may be of functional importance in the vision of frogs. For example, site 169 in *RH1* was found to be under positive selection in 12 lineages. Experiments using site-directed mutagenesis could help verify which sites are important for spectral tuning in amphibians.

In Table 1 we indicate that many sites are close to known spectral tuning sites, and may also act as spectral tuning sites themselves or could be under selection due to epistatic interactions with nearby sites. For instance, sites 166 and 262 in *LWS* are close to known spectral tuning sites 164 and 261 (Escobar-Camacho et al. 2020); mutations at each or all of these sites can shift λ_max_ of *LWS* (Asenjo et al. 1994;Yokoyama and Yokoyama 1990; Yokoyama and Radlwimmer 1998). Similarly, changes at site 50 or adjacent sites in *SWS1* could affect the spectral tuning site at position 49 and have an allosteric effect on *SWS1* sensitivity. Although amino acid replacements at site 49 do not shift the maximum wavelength by themselves, when they occur in conjunction with other substitutions, a shift in sensitivity occurs in mouse *SWS1* (Yokoyama & Tada 2003). Additionally, MEME and FEL identified positive selection in site 120 of *SWS1*, which is also reported by Schott et al. (2022) to be under positive selection. A mutagenesis experiment showed that a nearby site (position 118) is among the seven sites that shift λ_max_ of *SWS1* and affect violet sensitivity (Takahashi and Yokoyama 2005). Therefore, Table 1 provides an extensive list of sites putatively involved in frog vision. Changes at these sites may work alone or together to shift the wavelength sensitivity of anurans as they adapt to diverse light environments, including diurnality.

Of the 16 sites identified by Schott and colleagues (2022) to be under positive selection in opsin genes of anurans, we found 11 to be under positive selection in our study (Table 1). We fail to identify five sites (65, 212, and 270 for *RH1;* 154 for *LWS*; −2 for *SWS2*), which might reflect the differences in the number of taxa (i.e., 116 in our study vs. 33 in theirs) or the specific focus on diurnal species in our study. Schott et al. (2022) also discussed additional variation in other known spectral tuning sites. We exclude them from our discussion because we could not find evidence of positive selection on these sites. Our comparison depicts the conserved nature of opsins, but also reports new amino acid changes that contribute to the repertoire of variants that might contribute to opsin spectral sensitivity.

Initially, we aimed to identify sites and patterns of selection that were associated with the transition to diurnality in the three focal anuran clades sampled for this study (Dendrobatidae, *Atelopus* and *Brachycephalus*). However, our MEME results uncovered branch-site combinations that did not cluster on the nocturnal-to-diurnal transition branches (e.g., stem branches of clades that evolved diurnality independently; see Fig. 1). Moreover, the Contrast-FEL analysis, which we employed to detect sites under selection in diurnal lineages, revealed a bias in our sampling where sites associated with diurnality overlapped with sites under selection within the Dendrobatidae family.

Nevertheless, we were able to parse apart some of the patterns among Dendrobatidae and other diurnal lineages and propose the following. Seven sites (176 in *LWS*, 218 and 221 in *SWS1*, and 87, 124, 169, and 277 in *RH1*) were found to be under positive selection in diurnal lineages including Dendrobatidae, and two sites (318 in *LWS* and 22 in *RH1*) were found to be under positive selection in diurnal lineages excluding Dendrobatidae; these 9 sites appear to be associated with adaptation to diurnality. In contrast, site 213 in *RH1* was found to be under negative selection in diurnal lineages but under positive selection in nocturnal or crepuscular lineages, suggesting it might be important instead for dim-light vision. Finally, three sites show Dendrobatidae-specific selection patterns. Sites 316 in *LWS* and 336 in *SWS1* are under positive selection in Dendrobatidae but under negative selection in other branches; site 97 in *RH1* is under positive selection in all branches except dendrobatids, where it is under negative selection. We also speculate that the transition to diurnality was followed by diversification in visual opsin genes, and indeed we identify many sites under selection within the three diurnal lineages on which we focused (i.e., poison frogs, *Atelopus* and *Brachycephalus*), even if these were not found to be specifically associated with diurnality in the Contrast-FEL assessments. As many diurnal animals rely on color-based signals (i.e., detecting ripe fruits; [Melin et al. 2009]) or perceiving sexual signals ([Carleton et al. 2005]), they are expected to have a suite of opsins that are tuned to maximally perceive the relevant signal under the specific light environments that are present in the habitats where they live. In summary, our selection results suggest that the transition to diurnality is complex and that each lineage accumulated some changes to accommodate the diel habit. In other words, the opsin genes in each clade experienced the process of visual tuning independently; not surprisingly, this resulted in some parallel and some different substitution patterns.

### *Loss of* SWS2 *in Dendrobatidae and the implications for their visual ecology*

Based on our inability to sequence *SWS2* from an *O. pumilio* eye transcriptome, the complete absence of *SWS2* from bait-capture data from all major lineages of Dendrobatidae (60 species), and our failure to identify any trace of an *SWS2* gene or pseudogene in the *O. pumilio* and *R. imitator* genomes, we hypothesize that dendrobatids lost this gene early in their history. Such gene-loss events are not surprising as other comparative genomic studies have shown (Borges et al. 2015; Xu et al. 2021). The loss of short-wavelength photopigment genes has occurred multiple times in the evolutionary history of vertebrates and coincides with shifts in activity patterns, habitat occupancy, and the evolution of other aspects of the sensory capacity of animals (Bowmaker 2008; Jacobs 2013). For example, the loss of *SWS2* in therian mammals and coelocanths is considered an adaptation to nocturnality and deep sea environments, respectively. Whereas, the loss of a functional *SWS1* pigment in several bat species coincides with the origin of specialized form of echolocation (i.e. high-duty-cycle echolocation; Jacobs 2013).

In dendrobatids, it remains unclear how the loss of *SWS2* may have impacted their vision and visual ecology. To answer such a general problem, we can explore some specific questions. First, how has *SWS2* loss contributed to the differentiation of dendrobatids? The limited empirical data from anurans suggest that *SWS2* expression is restricted to green rods (Yovanovich et al. 2017; Mohun and Davies 2019). However, green rods are absent from the *O. pumilio* retina (Siddiqi et al. 2004). Thus, the loss of *SWS2* suggests that under low light levels, dendrobatids might be less capable of discriminating color (scotopic vision) than other diurnal frogs. To our knowledge, color vision and its molecular underpinnings in the other two focal clades of this study, *Atelopus* and *Brachycephalus*, remain unstudied. Our data suggest that only dendrobatids lost *SWS2*. Additional data may show that other groups (e.g., hylids) have lost *SWS2*, but until then, we consider the loss of *SWS2* is a synapomorphy of Dendrobatidae (i.e., Aromobatidae + Dendrobatidae sensu Grant et al. [2006]).

Second, how did *SWS2* disappear from the genome of ancestral dendrobatids? Based on our results, we could not find a *SWS2* pseudogene (i.e., nonfunctional segments of DNA that resemble functional *SWS2*) in the available genomic data, while close relatives of dendrobatids do have a functional *SWS2* (Fig. 2). Hyloidea is a superfamily of frogs that contains dendrobatids; the last ancestor of hyloids existed about ∼70 Mya (Feng et al. 2017; Hime et al. 2022). The approximate age of the crown clade of Dendrobatidae is ∼40 Mya (Feng et al. 2017) and thus *SWS2* must have been lost between 40–70 Mya. Two alternative explanations exist for the loss of *SWS2* during this 30-My timeframe: (1) The ancestor of dendrobatids had a functional gene which pseudogenized and changed beyond recognition during that period, or (2) *SWS2* was lost without pseudogenization. Given the lack of evidence of *SWS2* pseudogene relics, the most parsimonious explanation is a complete deletion of the gene.

Third, was the loss of *SWS2* consequential in that it affected other aspects of dendrobatid vision? Other diurnal hyloid clades in our dataset have not lost *SWS2*: all *Atelopus* and *Brachycephalus* frogs maintain this gene. Whether these clades have the green rod cell type (which expresses *SWS2*) remains unknown. Nevertheless, dendrobatids are known to differ in at least two visual properties from close relatives. First, the lenses of some dendrobatid species (*Oophaga pumilio, Epipedobates tricolor, Dendrobates auratus, D. leucomelas, Allobates femoralis* and *Adelphobates castaneoticus*) transmit less short-wavelength light (lens ***λ***_t50_ 413–425 nm; except for *D. leucomelas* lens ***λ***_t50_ 326 nm) than those of bufonids (*Rhinella marina*, *R. icterica*, *R. ornata*, *Bufo bufo*, *Rhaebo guttatus*, *Sclerophis maculata*, and *Atelopus varius* lens ***λ***t50 331–365 nm) and brachycephalids (*Brachycephalus rotenbergae, Ischnocnema parva, I. henseli* lens ***λ***t50 314–356 nm) (Donner and Yovanovich 2020; Yovanovich et al. 2020; Thomas et al. 2022). Because the typical peak wavelength sensitivity of *SWS2* rods (∼430 nm; Yovanovich et al. 2017) is much closer to the limit of lens transmission in dendrobatids than other frogs, it is plausible that lens transmission properties were adjusted in some species following the loss of *SWS2.* Additionally, in *O. pumilio*, the short-wavelength-sensitive cone (likely *SWS1*), which absorbs light wavelengths around 430 nm in other anurans (Yovanovich et al. 2017), was found to absorb light at 466 nm (Siddiqi et al. 2004), suggesting that it has undergone a shift in spectral sensitivity. Further investigation of these patterns will clarify whether loss of *SWS2* in dendrobatids in combination with a diurnal lifestyle led to altered lens transmission properties and a change in *SWS1* wavelength sensitivity, or vice versa.

Despite the apparent loss of *SWS2* in dendrobatid frogs and the lack of *RH2* pigment in amphibians (Mohun and Davies 2019), microspectrophotometry data revealed an *MWS* photoreceptor in the *O. pumilio* retina (Siddiqi et al. 2004). In accordance with prior literature (Schott et al. 2022), we speculate that these frogs are instead using *RH1* in those photoreceptors as the microspectrophotometry absorbance graphs for rods and MWS cones appear nearly identical (Siddiqi et al. 2004). Whether this is the case and if there are any modifications to the *RH1* pigment or to any proteins in the *O. pumilio MWS* cone phototransduction pathway required to make the rod protein function in cone cells requires further investigation. We did find that a large number of amino acid sites (i.e., 20) are under selection in *RH1* within frogs, which could be a result of this dual function, though only a few of these sites were noted to be specifically under selection in Dendrobatidae or experiencing dendrobatid-specific patterns of selection. Further, our genome mining, transcriptome, and synteny analyses do not provide any evidence suggesting that *RH1* has been duplicated in frogs, but at least one species (*Pyxicephalus adspersus*) has a duplication of *LWS* (Schott et al. 2022). Empirical data of spectral sensitivity and opsin protein function in frogs is sparse and further studies using microspectrophotometry of isolated poison frog rods and cones will be necessary to recapitulate the evolution of vision in frogs (Donner and Yovanovich 2020).

### Conclusion

Our results fill gaps in our knowledge by illustrating how diversification in ecology and life history may have affected opsin evolution in amphibians. We uncover evidence of new putative tuning sites in frogs and show strong evidence suggesting that poison frogs have lost the *SWS2* gene. Further work is needed to elucidate the functional consequences of its loss, and of the potential role of *RH1* in permitting nocturnal color vision in some frogs. Like Donner and Yovanovich (2020), we expect that additional studies of opsin evolution in amphibians will reveal non-canonical visual adaptations and broaden our understanding of the many ways in which animals adapt to diverse light environments.

## Materials and Methods

### Bait capture design, library preparation, and sequencing

Using publicly available sequences (see Table S1), previously generated transcriptomes (Santos et al. 2018), and sequences from a frog transcriptome project described in Santos et al. (2018), we designed a custom bait-capture array with myBaits® (Arbor Biosciences). The 120-bp baits were synthesized at >10x tiling across all exons of the genes of interest. Tissues from representative species of Dendrobatidae, *Atelopus*, and *Brachycephalus* and their relatives were obtained from the field or from museum collections (see Table S1 for source information), under approved protocols (UT Austin AUP-2012-00032 and AUP-0709210, STRI 200715122207, 2015-00205 Tulane #0453) and collection permits (001-13 IC-FAU-DNB/MA and 001-11 IC-FAU-DNB/MA [Ecuador], IBD0359 Res 1177-2014 and Isla Gorgona PIDB DTPA 020 - 16 Res 061-16 [Colombia], and SE/A-47-07 and CITES export permit SE/A-47-07 [Panama]). Tissue samples from Ecuador and Peru were loaned by Luis A. Coloma (Fundación Otonga and Centro Jambatu de Investigación y Conservación de Anfibios; Ecuador) and César Aguilar (Universidad Nacional Mayor de San Marcos; Lima, Peru). We note there controversy over the elevation of subgenera such as *Oophaga* to genera (Grant et al., 2017; Santos et al., 2009), but we have followed taxonomy following (AmphibiaWeb, 2023).

DNA was extracted using Qiagen DNeasy kits (Germantown, MD). DNA quality was reviewed with a 0.8% electrophoresis gel and highly degraded samples were excluded. RNA was removed from extractions with RNAse A (E1008, Zymo Research, Irvine, CA), and extractions were further purified and concentrated with Genomic DNA Clean and Concentrate (D4011, Zymo Research) and then quantified using a Qubit 3.0 fluorometer (Thermo Fisher Scientific, Waltham, MA) following manufacturer protocols. Given the large size of amphibian genomes, we used 400–1600 ng of starting DNA for each sample for library preparation. DNA was sheared to ∼300bp using a Covaris S2 Focused-ultrasonicator (Covaris, Inc, Woburn, MA; settings as Intensity: 5; Duty Cycle: 10%; Cycles per Burst: 200; Time: 50s; Temp: 7C; Water Level: 12; Sample Volume: 50 µL). Whole-genome libraries were prepared from sheared samples using the KAPA Hyper Plus library preparation kit (KK8514, Roche Diagnostics, Santa Clara, California), NEBNext Multiplex Oligos for Illumina (E7600, New England BioLabs, Ipswich, MA), home-made SPRI beads (Rohland and Reich 2012) (pers. comm. Lydia Smith), and manufacturer protocols. Uniquely barcoded libraries from 4–10 closely related species were pooled to a total of 1.6– 5µg DNA and then size-selected for an insert size of 250 ± 25 bp (total length with 120-bp adaptors: 345 ± 25 bp) using a Blue Pippin (Sage Science, Beverly, MA) and 2% gel cassette. For samples with low concentrations, a second batch of libraries were constructed with the same protocol, except that these were size-selected to 250 ± 50 bp with a 1.5% gel cassette (total length with 120-bp adaptors: 370 ± 50 bp). All size-selected libraries were cleaned with Qiagen MinElute PCR purification kits, eluted in 10 µL, and quantified with Qubit 3.0.

7µL of each pooled library set (50–275ng/pool) was then hybridized with custom myBaits biotinylated RNA, C0t-1 DNA, and xGen® Universal Blockers–TS Mix (1075474, Integrated DNA Technologies, Redwood City, CA) according to myBaits v4.01 recommended protocols. Because amphibian genomes are large and contain many repetitive sequences, we used a large amount of starting DNA (1.6–5µg at size selection) and a larger quantity of blocking oligos than described in the manufacturer’s protocol (8ug human C0t-1 and 8µg salmon C0t-1 per hybridization reaction). We also used the xGen blockers rather than those provided in the myBaits kit. Following hybridization with C0t-1, universal blockers, and baits for 36 h, pooled libraries were washed and cleaned with DynaBeads MyOne Streptavidin C1 beads (65002, Thermo Fisher Scientific). 12–17µL of resulting libraries were amplified using the NEBNext Illumina primers (that came with the multiplex oligos) and the KAPA Library Amplification Kit (KK2611, Roche Diagnostics) in two separate PCR reactions with 15-17 cycles. These were purified and eluted in 10µL with the Qiagen MinElute purification kit. Duplicate PCR reactions and libraries from all 97 samples were normalized by concentration and according to the number of samples per pool and then sequenced across two lanes of the Illumina HiSeq4000 at the Genome Sequencing and Analysis Facility (GSAF) at University of Texas at Austin, yielding approximately 4M reads per sample.

### Assembly of opsin genes

Read quality was checked with FastQC (Andrews 2010), barcodes were excluded with Trimmomatic (Bolger et al. 2014) and, because combining multiple assembly methods better characterize multi-gene families (Holding et al. 2018), the reads were assembled with default parameters for *de novo* with MEGAHIT (Li et al. 2015), Trinity (Grabherr et al. 2011) and SPAdes (Bankevich et al. 2012). Final assemblies were reduced using CD-HIT set to >98% similarity. Reduced assemblies were annotated using a custom library generated with the opsin sequences used to design the baits and BLASTX. Reference sequences for BLASTX were derived from amphibian opsin genes available in GenBank: *Xenopus laevis* (Pipidae), *Nanorana parkeri* (Dicroglossidae) and other species with opsin sequences (e.g., *Rana catesbeianus* [Ranidae]). Any sequence matching one of the reference opsin genes with an e-value <10^-6^ was pulled out for downstream analyses.

Given that we *de novo* assembled genomic DNA that was captured using baits designed from mRNA, the sequences were often only partially assembled, and some shorter sequences were identified erroneously by BLASTX. We used the program BLAT v.36×2 (Kent 2002) with reference sequences from the *Nanorana parkeri* genome (*LWS*, XM_018560714.1; *SWS1*, NW_017306744.1; *SWS2*, NW_017307939.1; *RH1,* NW_017306456.1) to verify sequences from our dataset that were putatively identified as opsins using BLASTX as described above. A BLAT server was prepared using *N. parkeri* reference sequences with the "-trans" option to translate the database into protein for all six reading frames. Then BLAT was run with options "-t=dnax" and "-q=dnax" to specify that the format of the database and query were both DNA sequences translated in six frames to protein. Sequences that matched *N. parkeri* references were pulled out from the BLAT results file generated using "-o=pslx". These query sequences were then aligned to the reference genome using MAFFT v47.19 (Katoh and Standley 2013) with the options "--auto" and "--adjustdirection". The smaller fragments whose sequences did not align to the exons were manually removed, and exons from the same individual of each species were merged.

For most of these genes, some edges of exons and several short exons were still missing from the assemblies. To recover the sequences or fill these gaps, we implemented an *in silico* target capture using MITObim (Hahn et al. 2013). For this procedure, we used the recovered exon sequences as bait sequences and the raw exome capture as target data. We included the following parameters "--quick" (starts process with initial baiting using provided fasta reference), "--mismatch" (% number of allowed mismatches in mapping), and "--kbait" (set kmer for baiting stringency). For the last two parameters we used low stringency (12-17 and 10-15, respectively) to bait more raw reads beyond those obtained from standard assemblers. This approach allowed MITObim to progressively expand the extremes of the bait sequences beyond the original gene reconstructions. The resulting sequences were then aligned with the original sequence matrices for each gene and full sequences from NCBI references (e.g., *Xenopus*, *Nanorana*, and *Rana*). In most cases, these extended sequences overlapped and were merged with the original ones to recover the missing regions. Lastly, we generated a clean alignment containing only coding regions that we used for the subsequent analyses. Raw sequencing coverage for *SWS2* past position 389 was low (perhaps because probes designed for that region did not effectively pull-down exons compared to other probes), so we exclude the remaining sequence to avoid any potential errors in its reconstruction.

### Gene tree construction

To check for any sequence contaminants, misidentifications, or assembly errors, we estimated a gene tree for each opsin, partitioned by codon position (length of alignment divided by three, because three bases constitute to a codon), using raxml-ng v.0.9.0 (Kozlov et al. 2019), the GTR+G model, and 200 bootstrap replicates to assess support. We found several sequences of different species to be identical for each gene, although in each case the identical sequences were from closely related species. For *LWS*, there were two sets of identical sequences: *Atelopus glyphus* and *A. spurrelli; Brachycephalus boticario* and *B. olivaceus*. For *SWS1* and *SWS2*, there was one set of identical sequences for each gene: *Atelopus glyphus, A. limosus* and *A. varius* for *SWS1*; *Brachycephalus pernix* and *B. pombali* for *SWS2*. For *RH1*, we found two sets of identical species: *Brachycephalus pernix* and *B. pombali; Phyllobates aurotaenia* and *P. terribilis*. Because there are no genomes for these species, it was not possible to verify the sequences, so we conservatively excluded all identical sequence sets from the following analyses. Finally, to check for chimeras, we aligned all four gene sequences to each other using MAFFT (Katoh and Standley 2013) and estimated a gene tree using raxml-ng v.0.9.0 (Kozlov et al. 2019), the GTR+G model, and 1000 bootstrap replicates to assess support (with bootstopping enabled).

### Determination of diel habits

Data on time of activity was compiled from primary and secondary literature, including species descriptions, taxonomic revisions, and books (see Table S1 for references). See Supplementary Information for details on the activity states for each species used in our analysis and the associated sources. Activity patterns were categorized as diurnal or nocturnal, with the latter category conservatively including species with exclusively crepuscular or mixed activity patterns. When not clearly stated, the time of activity was inferred from reports on time of calling, breeding activity, and behavior (asleep or active) at the time of collection. Diurnal groups include all of Dendrobatidae, *Atelopus*, *and Brachycephalus* (and the stem branches leading to each of these clades), as well as *Mantella baroni*, *Mantidactylus betsileanus*, and *Melanophryniscus stelzneri*. The following genera were classified as nocturnal: *Adenomera, Agalychnis, Amazophrynella, Amietia, Bufotes, Centrolene, Ceratophrys, Cochranella, Craugastor, Discoglossus, Fejervarya, Hyla, Hymenochirus, Ischnocnema, Leptobrachium, Limnodynastes, Lithodytes, Microhyla, Nanorana, Odorrana, Oreolalax, Osornophryne, Pelobates, Pyxicephalus, Quasipaa, Rana, Rhinella, Scaphiopus, Telmatobius,* and *Xenopus*.

### Analyses of selection

An updated phylogeny of the focal taxa was derived from alignments provided in the two largest phylogenetic reconstructions of amphibians (Pyron 2014; Jetz and Pyron 2018). Both alignments were appended, and taxa not included in our analysis were removed, duplicate taxa were also removed by choosing the one with more sequence data, and taxonomic nomenclature was updated following (AmphibiaWeb 2022). The final sequence alignment was realigned with MAFFT (Katoh and Standley 2013), then reviewed and adjusted manually, particularly the mitochondrial ribosomal gene sequences. Multiple alignment programs provide a good starting point, but they usually need to be examined and adjusted by eye (Baum and Smith 2013). The optimized alignment was then used for phylogenetic estimation. A maximum likelihood tree was estimated using IQ-TREE2 2.1.3 (Minh et al. 2020) with five replicate runs. The tree was constrained so that the topology among the families of Hyloidea matched that found by Feng et al. (2017) and Hime et al. (2020). This method was preferred because the trees presented in Pyron (2014) and Jetz and Pyron (2018) are largely dictated by the abundance of data from mitochondrial genes. In contrast, the trees found by Feng et al. (2017) and Hime et al. (2020) are based on 100-1000s of nuclear genes and no mitochondrial data. The sequences were partitioned by gene and codon position and the best-partitioned model was determined using - TESTMERGE option (Chernomor et al. 2016; Kalyaanamoorthy et al. 2017) within IQTREE2. Ultrafast bootstrap values (Hoang et al. 2018) were calculated using 50000 replicates and were plotted on the best likelihood tree. The data matrix, IQTREE2 scripts, constraint tree, and analysis log files are included in Supplementary Materials Data S3 in the folder IQTree.

For each selection analysis, we pruned the tree using the ‘ape’ R-package (Paradis and Schliep 2019) to contain the subset of species available for each opsin gene. In some cases, we replaced a tip in the tree with a closely related species that was used in our study (see Table S1 for details). We then used the resulting tree as the backbone for three types of site-based selection analyses (with p-value set to 0.05) in HyPhy version 2.5.14 (Kosakovsky Pond et al. 2005): FEL, which estimates the rate of synonymous (*dS*) and non-synonymous (*dN*) substitutions per site with maximum likelihood and compares them using likelihood ratio tests (Kosakovsky Pond and Frost 2005); FUBAR, a hierarchical Bayesian method that estimates diversifying positive selection by using a large number of predefined site classes (Murrell et al. 2013); MEME, a mixed-effects maximum likelihood approach, to test for positive selection using a branch model (Murrell et al. 2012); and Contrast-FEL, which compares substitution rates among sets of branches and sites to detect positive selection (Kosakovsky Pond et al. 2020). We created three sets of foreground lineages to compare with background lineages in Contrast-FEL: “DENDRO”, i.e., all Dendrobatidae plus its stem branch, “DIURNAL”, i.e., all diurnal branches as described above, and “DIURN-DEN”, which is all diurnal branches except for the Dendrobatidae lineages and its stem branch. Because there were no *SWS2* sequences for Dendrobatidae, we did not include this gene in Contrast-FEL analyses. To compare our results to those of Schott et al. (2022) we also conducted CODEML analyses comparing the M7 and M8 models (to detect positive selection on specific sites) in the PAML software (Yang 1997; Yang 2007). The type of selection is determined by the value of omega, which is calculated by rate of non-synonymous substitutions (dN) divided by rate of synonymous substitutions (dS). Omega value less than 1 indicates negative selection, while neutral selection is indicated by omega value equals to 1, and positive selection is depicted by omega value greater than 1.

Following convention, the amino acid sites are numbered based on the bovine rhodopsin sequence (NP_001014890.1); sequences from each opsin gene were aligned with bovine rhodopsin using MAFFT v7.419, and the numbering of the amino acid site was determined by referring to the numbering of bovine rhodopsin starting with the start codon as 1 (see Data S1). We report our selection results referring to the location of each amino acid according to the three-dimensional structure of opsins, which encompasses seven transmembrane domains (TMD I–VII), three extracellular domains (ECD I–III), and the amino-and carboxyl-termini (N and C) (Palczewski et al. 2000; Table 1, Fig. 1).

*Attempts to find* SWS2 *in* Oophaga pumilio *and* Ranitomeya imitator. Because we did not recover *SWS2* sequences from any dendrobatid species in our bait-capture dataset, we attempted to verify whether the *SWS2* gene had been lost in this clade using transcriptomics and genome skimming.

Transcriptomics-based approaches that include *de novo* transcriptome sequencing of *O. pumilio.* First, we attempted to determine which opsin genes are expressed in eye tissue of *O. pumilio.* As part of another project, one eye from each of eleven *O. pumilio* populations (Aguacate, Almirante, Bastimentos “Cemetery”, Bastimentos “Green”, Bastimentos “Orange”, Cayo Agua, Colón, Pastores, Popa, San Cristóbal, Solarte; see Maan and Cummings (2012) for details) was taken out of RNAlater and placed immediately in Trizol (Life Technologies, Grand Island, NY). RNA was extracted according to manufacturer instructions. Equal concentrations of total RNA from each of the 11 samples were pooled into one sample from which poly-adenylated RNA was isolated using the Poly(A) purist kit (Life Technologies) and manufacturer instructions. Lack of contaminating rRNA was confirmed using an Agilent 2100 Bioanalyzer. Strand-specific libraries for 100-bp paired-end sequencing were prepared and sequenced on the Illumina HiSeq 2000 according to manufacturer library kit instructions. A total of 83,168,029 reads were obtained. We preprocessed the reads using Trimmomatic (Bolger et al. 2014) by removing adapter sequence and sequence artifacts as well as trimming low-quality nucleotides based on the Phred score (Ewing et al. 1998) greater than 20, which corresponds to a 1% sequencing error rate.

*de novo* sequence assembly was then completed using Trinity (Grabherr et al. 2011) on the Odyssey cluster supported by the FAS Science Division Research Computing Group at Harvard University. We remapped reads to the raw assembly using BWA (Li and Durbin 2009) and then used eXpress (http://bio.math.berkeley.edu/eXpress/) to generate FPKM (fragments per kilobase per million mapped) scores for each contig. Low-confidence contigs that had an FPKM value of less than one were removed from the draft assembly. After removing contigs with low confidence, we used CD-HIT-EST (Li and Godzik 2006) to remove contig redundancy. Given that redundant contigs can represent alternative splice variants, polymorphisms among the pooled individuals, or sequencing errors, we used a conservative threshold of 98% sequence similarity. The final assembly was annotated using Trinotate (Bryant et al. 2017); it was found to contain only three opsin genes: *RH1*, *LWS*, and *SWS1*. The *SWS2* gene was absent. Thus, we hypothesized that *SWS2* might have been lost in the ancestor of dendrobatids.

Genomics-based approaches include a synteny analysis with *LWS* and mining of publicly available dendrobatid genomes. For the first approach, if the *SWS2* gene is no longer functional in dendrobatids, we predicted that remnants of *SWS2* might be detectable in genomic data as a pseudogene either near its expected location or translocated elsewhere in the genome. We thus reviewed the genomes (Rogers et al. 2018; Rodríguez et al. 2020) of *Oophaga pumilio* and *Ranitomeya imitator* (Stuckert et al. 2021) for evidence of *SWS2* using synteny and genome mining. We searched for the syntenic block containing *LWS* using Genomicus v.100.01 (Nguyen et al. 2018) with search term *opn1lw* and *Xenopus tropicalis* as the focal species. To visualize gene order in the syntenic block we used BLAST to locate *SWS2, LWS,* and the gene predicted to be directly upstream of *LWS*, mecp2 (methyl-CpG binding transcription factor) in *N. parkeri,* and *X. tropicalis* genomes. We hypothesized that a degraded ancestral *SWS2* sequence might be maintained in *O. pumilio* and *R. imitator* upstream of *LWS*. We mined the published genome assembly of *O. pumilio* (Rogers et al. 2018) and a re-scaffolded version of this genome (Rodríguez et al. 2020), as well as the recently published *R. imitator* genome for *LWS* using BLAST v2.10.0 (Altschul et al. 1990). We then attempted to align the scaffolds containing *LWS* against *SWS2* sequences from other amphibians using MAFFT v7.453 (Katoh and Standley 2013). We note that the *O. pumilio* scaffold contains two sections of Ns upstream of *LWS* (2093 and 2391 nucleotides long), which may inflate our perceived length of coverage of this chromosome. Likewise, it is possible that this scaffold might be misassembled and require further refinement and deeper coverage. The re-scaffolded version (Rodríguez et al. 2020) of the *O. pumilio* scaffold containing *LWS* is identical to the original and thus did not alter our conclusions. As a positive control for this analysis, we also ran our pipeline on the draft genome of *Rhinella marina*, which is known to contain *SWS2* based on our data. This analysis is implemented in the Github folder named “synteny” (see Data S4).

Lastly we also mined two dendrobatid genomes for *SWS2.* Because we were unable to detect *SWS2* upstream of the *O. pumilio* and *R. imitator LWS* scaffolds, we then hypothesized that the syntenic relationship between *LWS* and *SWS2* might have been broken in *O. pumilio* and *R. imitator*. Therefore, we used tblastn v2.10.0 to screen all scaffolds from the re-scaffolded genome assembly of *O. pumilio* and the draft assembly of *R. imitator* for any potential *SWS2* orthologs. This approach uncovered several candidate sequences with low E-values (<1e-10). We screened the top candidates using tblastn against *Nanorana parkeri*, a well-annotated frog genome more closely related to dendrobatids than *Xenopus*. None of the candidate sequences returned *SWS2* as the most likely ortholog; the highest scoring sequences were annotated as related proteins (e.g., pinopsin, rhodopsin; Table S4). As a positive control for this analysis, we also ran tblastn on 43 known *SWS2* sequences from other frog species against *Nanorana parkeri*. As *SWS2* is known to be present in *Nanorana*, we would expect our pipeline to correctly detect and annotate these sequences. We found that the top hit from *N. parkeri* correctly identified *SWS2* as the most likely ortholog for all 43 known *SWS2* sequences (Table S4). As another positive control, we ran an identical search for *LWS*, which identified *LWS* as the top hit for 101 other frog sequences and for one of the *Oophaga pumilio* scaffolds (Table S4). Lastly, we ran the reciprocal best-hit pipeline for *SWS2* using the *Rhinella marina* genome as the query. Since *Rhinella marina* is known to contain *SWS2,* we would expect that our pipeline should identify *SWS2* in the *R. marina* genome. We found that the pipeline identifies the same contig in *Rhinella marina* that is identified in our synteny-based analysis (ONZH01019223.1) as a likely *SWS2* ortholog, while other sequences with low E-values were annotated with other related opsin genes (Table S4). This analysis is implemented in the Github folders named “SWS2_search” and “LWS_search” (see Data S4).

## Acknowledgments

This research was funded by NSF DEB-1556967 to DCC; NSF GROW, NSF DDIG (DEB-1404409), National Geographic Society 2014 Young Explorer Grant, and UC Berkeley start-up funding to RDT; NIH 5P50GM068763, NSF IOS-1822025 and William F. Milton Fund from Harvard Medical School to LAO; LAO is a New York Stem Cell Foundation – Robertson Investigator. JCS was supported by SJU start-up funds and NSF DEB-2016372. MJN was supported by the American Association of University Women (AAUW). We thank Roberto Ibañez and Brian Gratwicke (Amphibian Rescue and Conservation Project), Luis A. Coloma (Fundación Otonga and Centro Jambatu de Investigación y Conservación de Anfibios; Quito, Ecuador), César Aguilar (Universidad Nacional Mayor de San Marcos; Lima, Peru), and Travis LaDuc (Biodiversity Center, UT Austin) for facilitating access to tissues and specimens. We thank two anonymous reviewers for critiques that improved the quality of this paper.

## Data accessibility statement

Raw Illumina sequencing data are deposited in the Sequence Read Archive (BioProject PRJNA856227). Specimens are available in QCAZ at Pontificia Universidad Católica del Ecuador, Centro Jambatu de Investigación y Conservación de Anfibios (Quito, Ecuador), Museo de Historia Natural de la Universidad Nacional Mayor de San Marcos (MUSM, Lima, Peru), Coleção de Herpetologia, Departamento de Zoologia, Universidade Federal do Paraná, (Paraná, Brazil), or Museo de Historia Natural ANDES at the Universidad de los Andes. All scripts and other data are available in the supporting material on DataDryad (DOI: 10.5061/dryad.zw3r2289j).

