## Supplementary material for "Selection on visual opsin genes in diurnal Neotropical frogs and loss of the *SWS2* opsin in poison frogs"

ORDER OF FIGURES ATTACHED TO THIS DOCUMENT (COMPLETE LEGENDS ARE PROVIDED IN-TEXT AND IN THE SUPPLEMENTARY MATERIAL DOCUMENT):

1. Figure 1. A summary of activity trait data used in analyses (see Table S1 for references) and results from HyPhy and CODEML selection analyses of two opsin genes, *LWS* and *RH1*.
2. Figure 2. A) SWS2 is found in a syntenic block with LWS in the ancestors of all tetrapods (block 355) and in the ancestor of all amniotes (block 65). B) Comparison of the syntenic block containing *SWS2*, *MECP2*, *IRAK1*, *TMEM187*, and *HCFC1* upstream of *LWS* among anuran species.
3. Figure S1. A summary of activity trait data used in analyses (see Table S1 for references) and results from HyPhy and CODEML selection analyses of two opsin genes, *SWS1* and *SWS2*.

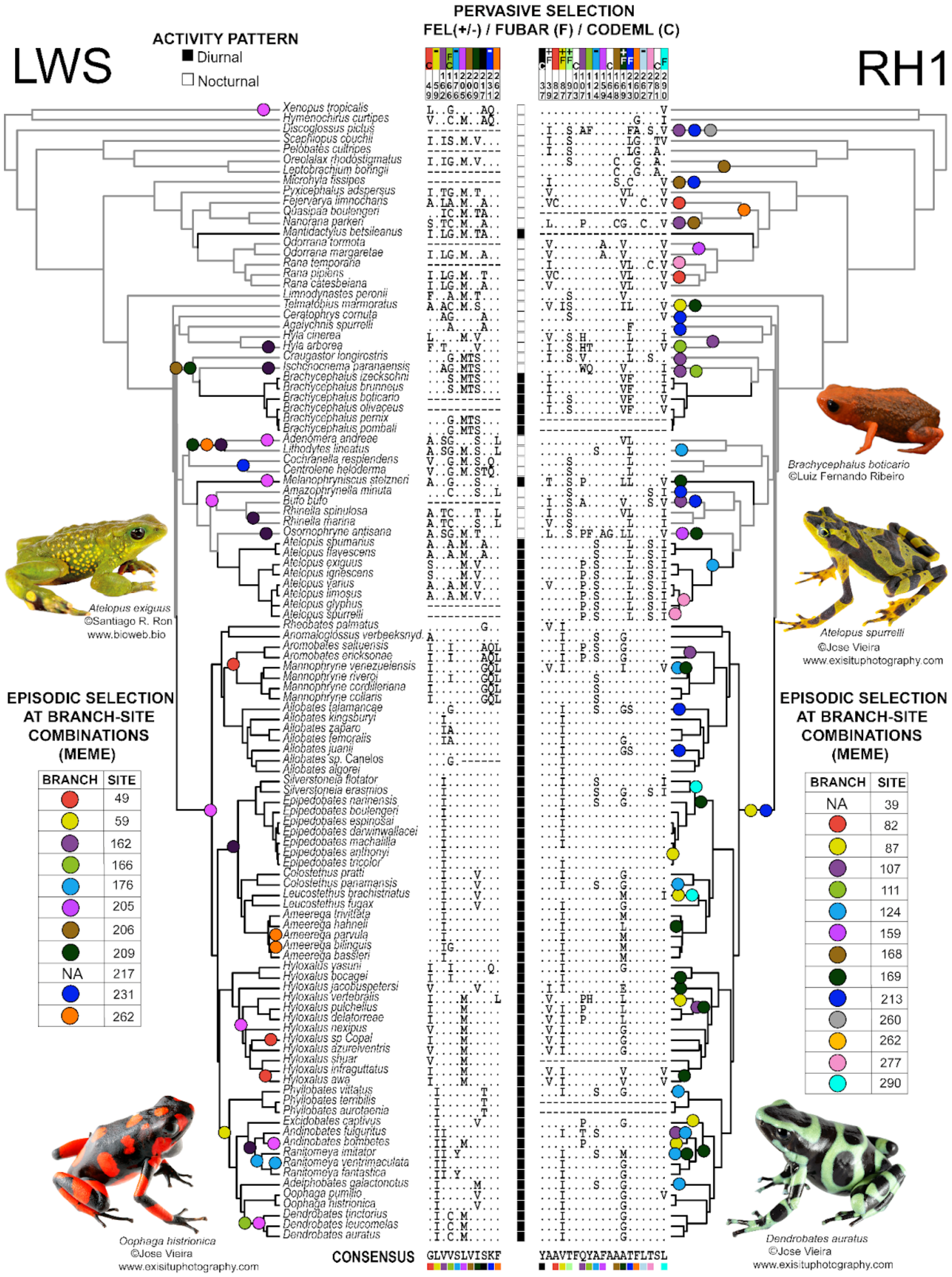

LWS
RH1
ACTIVITY PATTERN
Diurnal
Nocturnal
PERVASIVE SELECTION
FEL(+/-) / FUBAR (F) / CODEML (C)
Xenopus tropicalis
Hymenochirus curtipes
Discoglossus pictus
Scaphiopus couchii
Pelobates cultripes
Oreolalax rhodostigmatus
Leptobrachium boringii
Microhyla fissipes
Pyxicephalus adspersus
Fejervarya limnocharis
Quasipaa boulengeri
Nanorana parkeri
Mantidactylus betsileanus
Odorrana tormota
Odorrana margaretae
Rana temporaria
Rana pipiens
Rana catesbeiana
Limnodynastes peronii
Telmatobius marmoratus
Ceratophrys cornuta
Agalychnis spurrelli
Hyla cinerea
Hyla arborea
Craugastor longirostris
Ischnocnema paranaensis
Brachycephalus izecksohni
Brachycephalus brunneus
Brachycephalus boticario
Brachycephalus olivaceus
Brachycephalus pernix
Brachycephalus pombali
Adenomera andreae
Lithodytes lineatus
Cochranella resplendens
Centrolene heloderma
Melanophryniscus stelzneri
Amazophrynella minuta
Bufo bufo
Rhinella spinulosa
Rhinella marina
Osornophryne antisana
Atelopus spumarius
Atelopus flavescens
Atelopus exiguus
Atelopus ignescens
Atelopus varius
Atelopus limosus
Atelopus glyphus
Atelopus spurrelli
Rheobates palmatus
Anomaloglossus verbeeksnyd.
Aromobates saltuensis
Aromobates ericksonae
Mannophryne venezuelensis
Mannophryne riveroi
Mannophryne cordilleriana
Mannophryne collaris
Allobates talamancae
Allobates kingsburyi
Allobates zaparo
Allobates femoralis
Allobates juanii
Allobates sp. Canelos
Allobates algorei
Silverstoneia flotator
Silverstoneia erasmios
Epipedobates narinensis
Epipedobates boulengeri
Epipedobates espinosai
Epipedobates darwinwallacei
Epipedobates machalilla
Epipedobates anthonyi
Epipedobates tricolor
Colostethus pratti
Colostethus panamensis
Leucostethus brachistriatus
Leucostethus fugax
Ameerega trivittata
Ameerega hahneli
Ameerega parvula
Ameerega bilinguis
Ameerega bassleri
Hyloxalus yasuni
Hyloxalus bocagei
Hyloxalus jacobuspetersi
Hyloxalus vertebralis
Hyloxalus pulchellus
Hyloxalus delatorreae
Hyloxalus nexipus
Hyloxalus sp Copal
Hyloxalus azureiventris
Hyloxalus shuar
Hyloxalus infraguttatus
Hyloxalus awa
Phyllobates vittatus
Phyllobates terribilis
Phyllobates aurotaenia
Excidobates captivus
Andinobates fulguritus
Andinobates bombetes
Ranitomeya imitator
Ranitomeya ventrimaculata
Ranitomeya fantastica
Adelphobates galactonotus
Oophaga pumilio
Oophaga histrionica
Dendrobates tinctorius
Dendrobates leucomelas
Dendrobates auratus
CONSENSUS GLVVSLVISKF
YAAVTFQYAFAAATFLTSL
Brachycephalus boticario
©Luiz Fernando Ribeiro
Atelopus exiguus
©Santiago R. Ron
www.bioweb.bio
Atelopus spurrelli
©Jose Vieira
www.exisituphotography.com
EPISODIC SELECTION AT BRANCH-SITE COMBINATIONS (MEME)
BRANCH SITE
49
59
162
166
176
205
206
209
NA 217
231
262
EPISODIC SELECTION AT BRANCH-SITE COMBINATIONS (MEME)
BRANCH SITE
NA 39
82
87
107
111
124
159
168
169
213
260
262
277
290
Oophaga histrionica
©Jose Vieira
www.exisituphotography.com
Dendrobates auratus
©Jose Vieira
www.exisituphotography.com

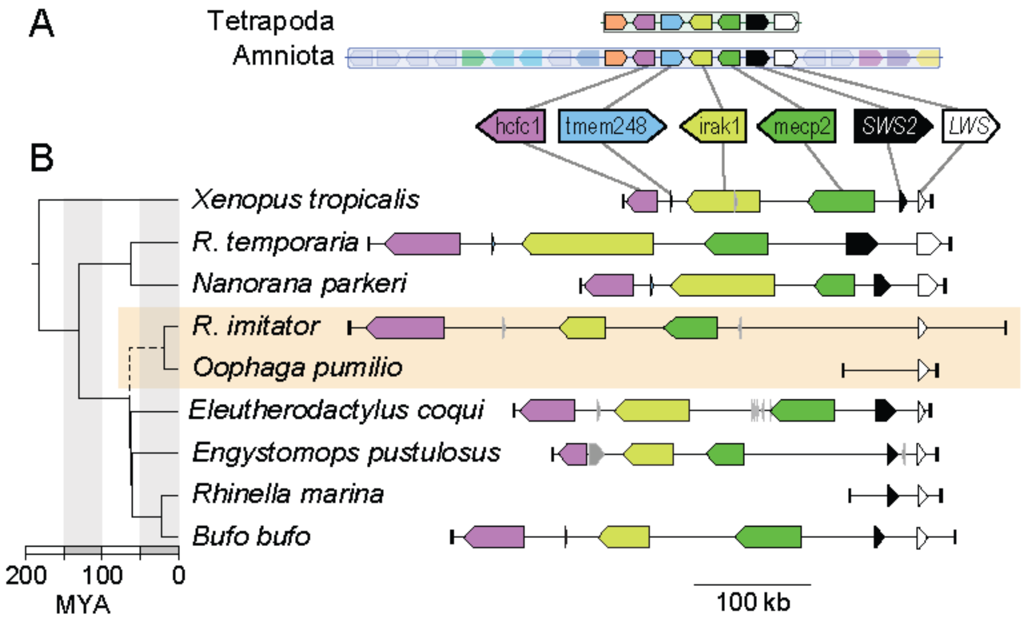
A
Tetrapoda
Amniota
hcfc1
tmem248
irak1
mecp2
SWS2
LWS
B
Xenopus tropicalis
R. temporaria
Nanorana parkeri
R. imitator
Oophaga pumilio
Eleutherodactylus coqui
Engystomops pustulosus
Rhinella marina
Bufo bufo
200
100
0
MYA
100 kb

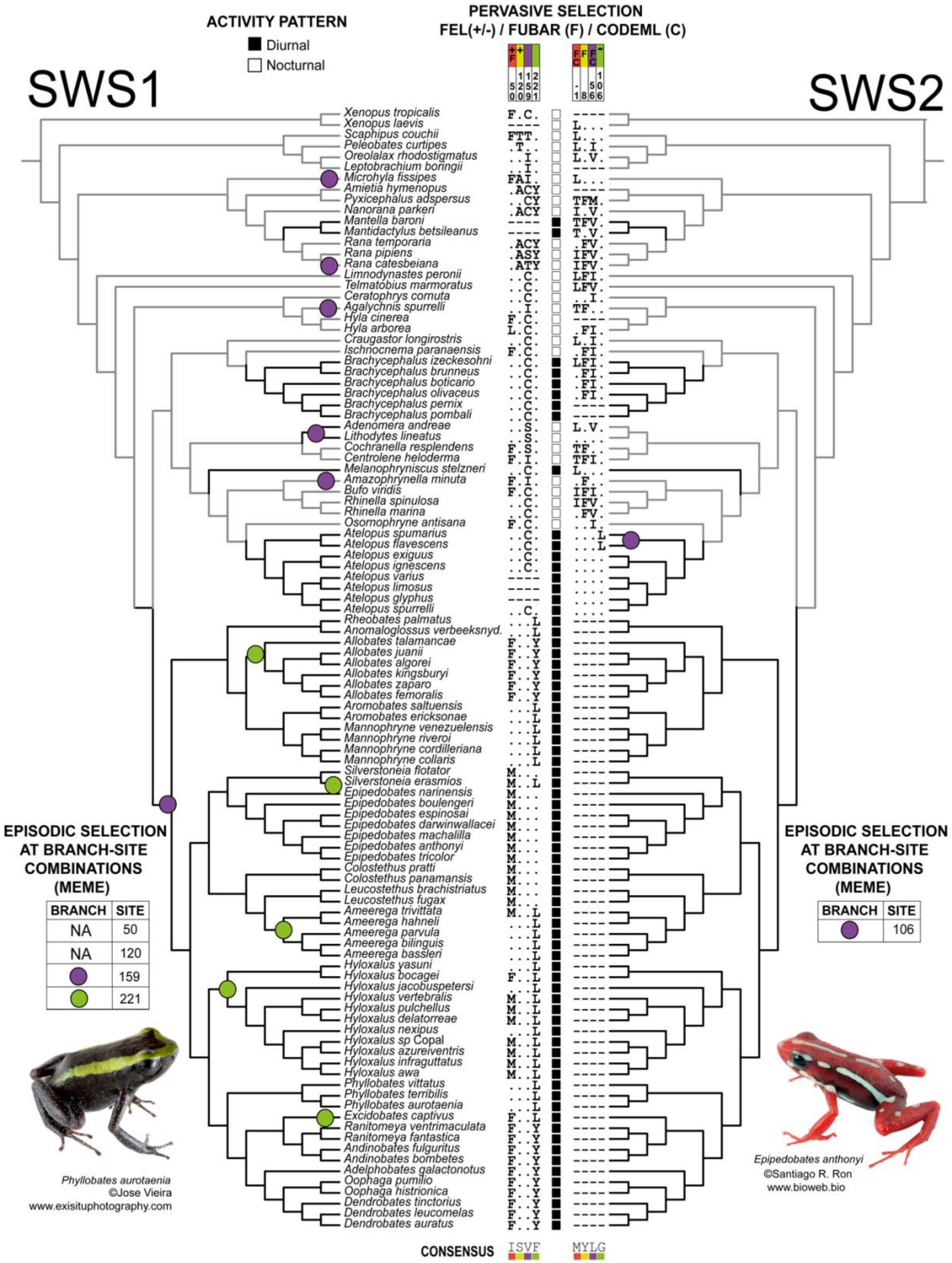

SWS1
SWS2
ACTIVITY PATTERN
Diurnal
Nocturnal
PERVASIVE SELECTION
FEL(+/-) / FUBAR (F) / CODEML (C)
Xenopus tropicalis
Xenopus laevis
Scaphipus couchii
Peleobates curtipes
Oreolalax rhodostigmatus
Leptobrachium boringii
Microhyla fissipes
Amietia hymenopus
Pyxicephalus adspersus
Nanorana parkeri
Mantella baroni
Mantidactylus betsileanus
Rana temporaria
Rana pipiens
Rana catesbeiana
Limnodynastes peronii
Telmatobius marmoratus
Ceratophrys cornuta
Agalychnis spurrelli
Hyla cinerea
Hyla arborea
Craugastor longirostris
Ischnocnema paranaensis
Brachycephalus izeckesohni
Brachycephalus brunneus
Brachycephalus boticario
Brachycephalus olivaceus
Brachycephalus pernix
Brachycephalus pombali
Adenomera andreae
Lithodytes lineatus
Cochranella resplendens
Centrolene heloderma
Melanophryniscus stelzneri
Amazophrynella minuta
Bufo viridis
Rhinella spinulosa
Rhinella marina
Osornophryne antisana
Atelopus spumarius
Atelopus flavescens
Atelopus exiguus
Atelopus ignescens
Atelopus varius
Atelopus limosus
Atelopus glyphus
Atelopus spurrelli
Rheobates palmatus
Anomaloglossus verbeeksnyd.
Allobates talamancae
Allobates juanii
Allobates algorei
Allobates kingsburyi
Allobates zaparo
Allobates femoralis
Aromobates saltuensis
Aromobates ericksonae
Mannophryne venezuelensis
Mannophryne riveroi
Mannophryne cordilleriana
Mannophryne collaris
Silverstoneia flotator
Silverstoneia erasmios
Epipedobates narinensis
Epipedobates boulengeri
Epipedobates espinosai
Epipedobates darwinwallacei
Epipedobates machalilla
Epipedobates anthonyi
Epipedobates tricolor
Colostethus pratti
Colostethus panamansis
Leucostethus brachistriatus
Leucostethus fugax
Ameerega trivittata
Ameerega hahneli
Ameerega parvula
Ameerega bilinguis
Ameerega bassleri
Hyloxalus yasuni
Hyloxalus bocagei
Hyloxalus jacobuspetersi
Hyloxalus vertebralis
Hyloxalus pulchellus
Hyloxalus delatorreae
Hyloxalus nexipus
Hyloxalus sp Copal
Hyloxalus azureiventris
Hyloxalus infraguttatus
Hyloxalus awa
Phyllobates vittatus
Phyllobates terribilis
Phyllobates aurotaenia
Excidobates captivus
Ranitomeya ventrimaculata
Ranitomeya fantastica
Andinobates fulguritus
Andinobates bombetes
Adelphobates galactonotus
Oophaga pumilio
Oophaga histrionica
Dendrobates tinctorius
Dendrobates leucomelas
Dendrobates auratus
CONSENSUS
ISVF
MYLG
EPISODIC SELECTION AT BRANCH-SITE COMBINATIONS (MEME)
BRANCH | SITE
NA | 50
NA | 120
(purple) | 159
(green) | 221
EPISODIC SELECTION AT BRANCH-SITE COMBINATIONS (MEME)
BRANCH | SITE
(purple) | 106
Phyllobates aurotaenia
©Jose Vieira
www.exisituphotography.com
Epipedobates anthonyi
©Santiago R. Ron
www.bioweb.bio
